# Simulation of the effect of the randomness of climbing fiber input on the relationship between motor learning and Focal Task-Specific Dystonia

**DOI:** 10.1101/2022.05.16.492217

**Authors:** Kaito Muramatsu, Shoko Yuki, Hiroshi Kori, Dai Yanagihara

## Abstract

Focal Task-Specific Dystonia (FTSD) is an intractable neurological disorder with no clear prevention or treatment that affects more than 1% of classical musicians and threatens the active lives of experts due to its task-specific tremor symptoms. In the present study, we focused on the motor learning function of the cerebellum, which has not been the focus of much attention in the past. We numerically simulated the firing of cerebellar Purkinje cells and cerebellar nuclei during eyeblink conditioning as a typical example of cerebellar-related timing motor learning, with the aim to find the principle of the pathogenesis of FTSD at the level of individual neurons. The results showed the sustained firing of cerebellar nuclei after the learning condition in which the climbing fiber input to Purkinje cells was continued randomly. Therefore, the present study claims a suggestive factor regarding the neural mechanism of the cerebellum in the motor learning-induced task-specific tremor, which is a symptom of FTSD. We also proposed a motor learning paradigm, “undesirable motor learning,” in which the motor goal is too advanced to be achieved by repetitions alone and converges to a different result than the desired.

## Introduction

Focal task-specific dystonia (FTSD) [1][2] is an intractable neurological disorder that affects experts in sports and more than 1% of classical music performers [3], while more less in not experts [4][5]. It presents with task-specific tremor symptoms that are lifethreatening to professionals. Previous research questionnaires and psychological experiments have identified “perfectionism” and “excessive repetitive practice” as factors in the development of FTSD [6], and some previous research has reported the relationship between FD and cerebellum [7] for instance “cerebellar hyperactivity” during seizures by using PET [8]. There have been several mice studies on dystonia and cerebellar cells; White et al. genetically knocked out Purkinje cell side receptors for climbing fiber system inputs, causing constant dystonic motor deficits [9], and Hisatsune et al. have reported that chemical manipulation of the climbing fiber system to induce an excess of climbing fiber system inputs that cause complex spikes to Purkinje cells (PKJ) causes dystonia-like tremors in mice [10]. Although both studies refer to the condition that is congenital and not task-specific, they are essential in considering the mechanism of dystonia-like muscle contraction in FTSD. Therefore, in this study, we focused on the neural mechanism of motor learning in the cerebellum, which has not received much attention as a pathogenic mechanism of FTSD and aimed to explore the pathogenic mechanism of acquired, FTSD-like seizures at the single-cell level. Motor learning generally consists of multiple types of information, including spatial and temporal information [11]. On the other hand, in investigating neural mechanisms, the relationship between information and outcome in motor learning can be clarified by narrowing down the information. In this respect, eyeblink conditioning [12] with features limited to temporal information has already been studied in various vertebrates [13], including humans [14], and simulations [15] at the level of neuronal mechanisms as cerebellum-related motor learning. Therefore, we used this paradigm to simulate the cerebellar cortical neuronal network in this study.

## Methods

### Motor learning model

In this study, we used eye-blink conditioning as motor timing learning, which consists of two variations; 1) conditioned stimulus (CS) was followed by an unconditioned stimulus (US) after 500ms intervals from CS-onset, which the time length of the CS-US (Inter-Stimulus Interval: ISI). 2) the ISI was normally distributed, at which σ= 100 *or* 200 ms. In the simulation, trials (1) or (2) were repeated 100 times.

### Network model

Yamazaki model [15][16], a cerebellar-dependent motor learning network structure in the previous study, defined the input from the pre-cerebellar nucleus (PN) and simulated eyeblink conditioning, which is a cerebellar-timed motor learning process, that can be reproduced by spatially arranging granule cells and Golgi cells in the network up to the output of the cerebellar nucleus. And Yamazaki showed that the dynamics of granule cells and Golgi cells in the recurrent network generate the expression of the passage of time by the transitions of collective granule cell cluster firings with a period of about 100 ms. In the present study, we performed simulations using the neuronal network shown in Figure 1 as an intercellular model that simplifies the Yamazaki model by directly defining inputs derived from parallel fibers (PF) and mossy fibers (MF). The intention is to preserve the cellular dynamics of temporal information in learning while facilitating the simulation of input signals, as described below. In a previous model [15], 16 PKJs receive the same US input from one IO and projected outputs to one CN. Based on the temporal representation of the CS showed already, it is sufficient for examining changes in the firing of individual CN in response to US randomization via PKJ cell that both PKJ, CN, and inferior olive nuclei (IO) are one each. The PKJ receives 11 parallel excitatory inputs from the PF cluster model. Each synapse between neurons is given an individual synaptic weight, and each of the 11 PF clusters is also given an individual synaptic weight. In addition, we did not define complex spikes in this study because we were testing how LTD generates short-term memory as synaptic plasticity through repeated motor learning trials. Therefore, only the synaptic plasticity due to LTD was calculated.

**Figure 1.**
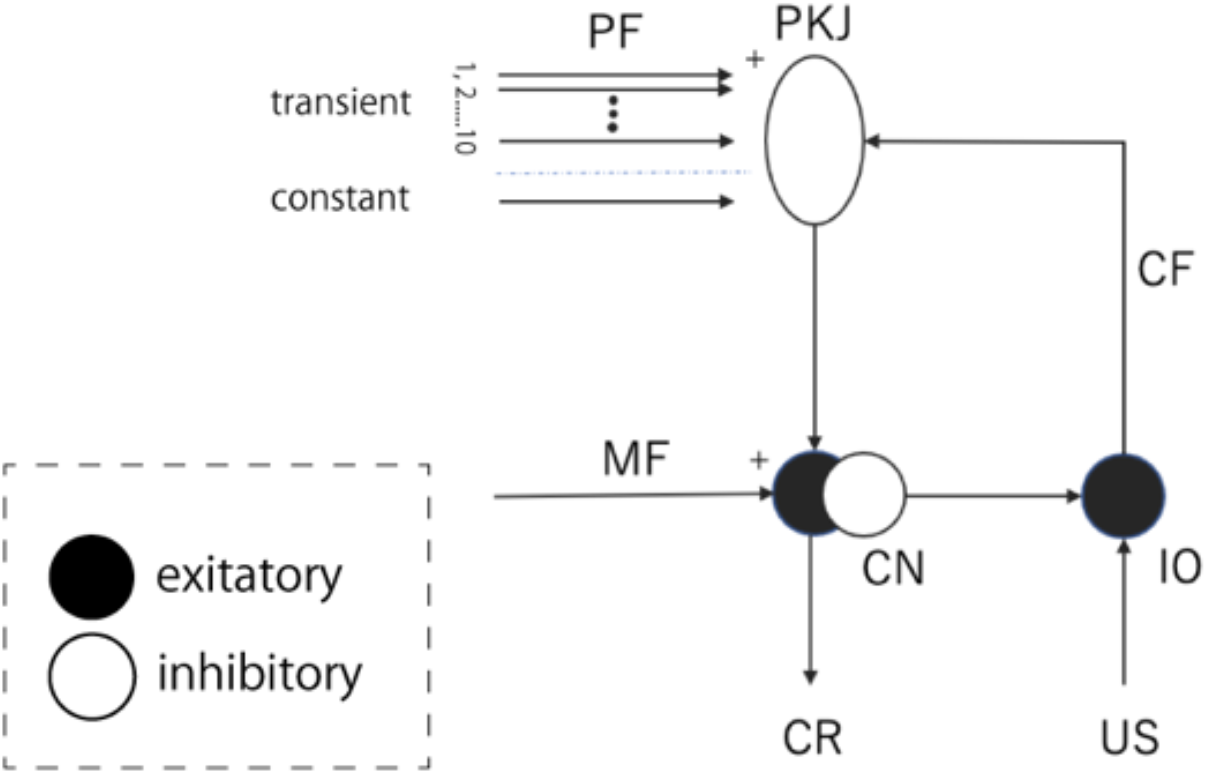
Schematic of cell types and synaptic connections incorporated into the present model with the flow diagram.

### Neural model

Leaky integrator model was used as the spiking model for PKJ and CN respectively expressed by the following equations:

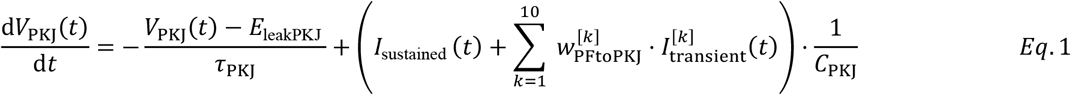

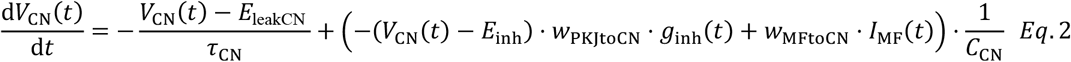

where *V*_PKJ_ and *V*_CN_ are the membrane potentials of PKJ and CN [mV], *E_leak_* is the reversal potential of each cell [mV], *E_inh_* is the reversal potential of inhibitory synapses [mV], *I* is the input current to each cell [pA], *C* is the cell capacitance [pF], *w* is the intersynaptic dimensionless coefficients as synaptic weights. The synaptic weights were adjusted to obtain conventional firing frequencies [15]. In addition, *τ* is the time constant [ms] expressed as follows:

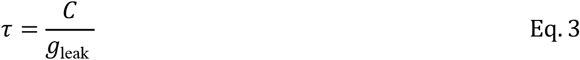

*where C* and *g*_leak_ are the capacitance [pF] and conductance [nS] determined for each cell, respectively. The inhibitory output from PKJ cells to the CN is expressed with conductance *g*_inh_,

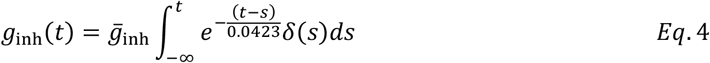

by substituting Eq.4 into Eq.1 and Eq.2. Here 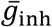 is the maximum conductance [nS], to which the alpha function 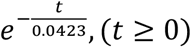 for the inhibitory input response of the CN and the delta function *δ*(*s*) with the spike time of PKJ as *s*[*s*] (0[s] ≤ *s* ≤ 3.0[s]) are convoluted. The specific values of each parameter followed that of previous studies [15].

### Stimulus

The current inputs to PKJ and CN defined in Eq.1 and Eq.2 correspond to the input stimuli from PF and MF, which are directly defined in this study. First, two types of currents were defined for the PF input to PKJ: constant and transition components. The constant component was defined as a 250 nA steady-state current with the intention of averaging the firing of various granule cell clusters, including clusters that do not fire at all and clusters that fire constantly [15]. Ten PFs were prepared for the transition component to represent the time course. The total current was 25 nA for the first and the last 1 second long. From CS_onset for 1 second, the current flowed 15 times the constant component on a different PF every 100 ms, and otherwise no current flow at all. Next, we defined the input signal from the MF to the CN. This has an overexcitability component that increases only during the presence of the CS and an unchanging constant component in the ratio of 1: 1 (i.e., 250nA) [17]. Finally, the climbing fiber input to PKJ was defined. Since only the learning effect was simulated this time, we did not define the complex spike, so we only simulated the LTD in the next section.

### Plasticity

We defined the LTD for PF-PKJ synapse within 5 ms before the time of the input to PKJ from PF as follows:

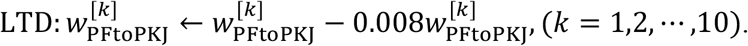

The rate of linear variation of LTD is a value that adjusts the intensity per exercise learning of the simulation [15]. As for LTP, it was defined by the following equation:

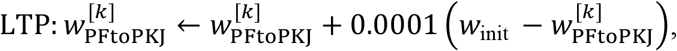

where *w*_init_ is the initial value of 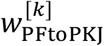 (i.e., 1).

### Simulation tool

The simulation program was created using python, and differential equations were calculated in 1 ms increments using the fourth order Runge-Kutta method.

## Results

Figure 2 shows the changes in CN membrane potentials during repeated CS-US paired presentations using the model described in the previous section. The membrane potentials in the CN increased right after the CS period and began to fire as trials progressed. When the ISI was fixed (σ= 0s), the CN began to fire after a few trials, and the time range of CN firing was limited to before the US presentation. When the ISI was randomized (σ= 0.1s, 0.2s), the CN continued to fire for a relatively long time regardless of the US input time of the trial. As shown in figure 3, interpreting a fixed ISI with standard deviation σ = 0s, as the standard deviation in the normal distribution increases, the number of trials in which the CN starts to fire increases and the time range in which the CN fires increases. In addition, the region at the time point it started firing once continued to fire in subsequent trials.

**figure 2.**
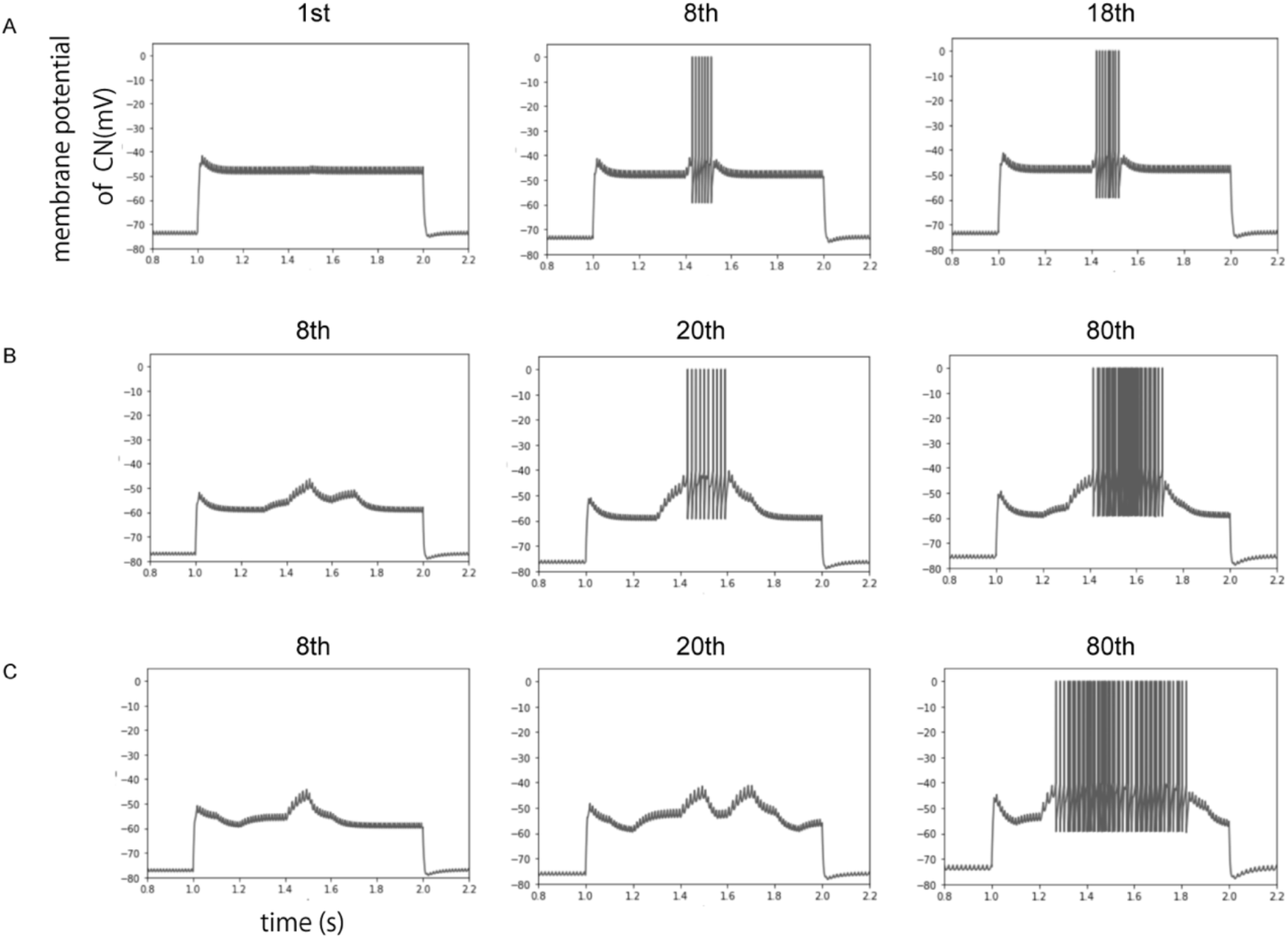
(A) Membrane potentials of CN at each trial (1st, 8th, 18th) in eyeblink conditioning with ISI fixed at 500 ms. (B)(C) Membrane potentials at each trial (8st, 20th, 80th) of CN in eyeblink conditioning with ISI rondomed at σ =100ms and 200ms respectively.

**Figure 3.**
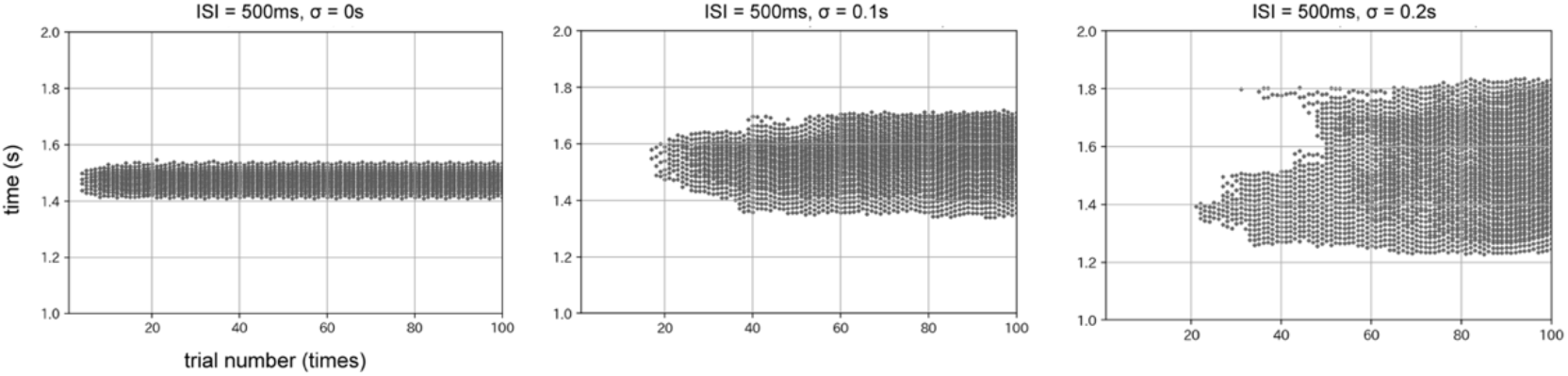
Plots of spiking of CN through trials, fixed ISI at 0.5s and random at σ = 0.1s and 0.2s, respectively.

## Discussion

### Simple time-related motor learning simulation model

The simplification of the previous model was done by the notion of time-related information in motor learning, as exemplified by eyeblink conditioning, which is represented by transitions of clusters of PF inputs [15]. The dynamics of granule cell clusters, which represent the passage of time from sound presentation (CS) in the eyeblink conditioning, are formed by a closed loop between granule cells and Golgi cells through the glomerulus and then sent to PF.

Therefore, we directly defined the PF input in the present model and simulated only the cellular dynamics after that. From the results of figure2-A, the firing dynamics of the CN appeared as eyeblink conditioning, which is a cerebellar motor timing learning, as in the previous study [13][14][15]. For these reasons, the abstracted representation of contextual temporal information as eyeblink conditioning is preserved from the prior model [15]. This model can simulate eyeblink conditioning and cerebellar-dependent timing learning.

### Undesirable motor learning results

The simulation results of the randomization of the ISI suggest that the CN may fire continuously over a wide range of time if the timing of the CF system input, which is the instructional signal in timing learning, is not fixed over sequential trials. This result suggests that LTD is induced at the PF-PKJ synapse for a broader period than in the conventional ISI-fixed eyeblink conditioning and that CN are more excitable for a longer period during CS. This is not considered to be a serious problem in the blink reflex, since the correct answer is to keep the eyelids closed (CN excited) during the US, but in conditions where the CN-onset and US-onset should be synchronized, such as the motion of hitting a ball, situations in which an error of tens hundreds of ms can produce a significant difference in results. If imposed, this may be an undesired motor learning outcome.

### Relationship between FTSD and undesired motor learning

In previous studies of FTSD, attention has gradually focused on the cerebellar origin contribution of symptoms [4][5]. In pharmacological and genetic manipulation studies in animals, manipulations on PKJ and CN have been able to induce dystonia-like spasms while constant. The gap between these studies and the symptoms of FTSD, the mechanism by which cerebellar-derived FTSD symptoms can also be caused by motor learning alone, the results of this study provide a prediction for this, while also ensuring that the ataxia is motor specific. In other words, the continued uncertain sequence of CF input timing, which is error information in repetitive motor learning, can be explained as a situation of repeated undesired motor learning, as defined in the previous section. And as a result, it may cause long excitation of the CN triggered by the CS at least in timing learning (i.e., loss of precise and advanced timing learning), and the uncertainty of the trial-by-trial teacher input may be a principle/influencing factor in the onset of FTSD from a neural circuit perspective. It has also been suggested that contextual information of any movement, not only timing learning, can be represented by the collective firing of PFs. [18][19] Therefore, the possibilities presented in this study are not limited to timing learning alone but are intended to be applied in a complex way of learning, including gain learning, etc.

### limitations

In the present simulations, signals from PF and MF were directly defined, and the climbing fiber system input was also assumed to always be provided in response to unconditioned stimuli. While we described the potential risk of overlearning in the cerebellum using an abstracted model, in actual motor learning in living organisms, all kinds of motor context information, including not only timing information but also gain information, are input. Therefore, to make the quantitative risk of FTSD in vivo applicable based on the present findings, it is at least necessary to conduct actual experiments of motor learning in mice, such as eyeblink conditioning learning as in the present study. By verifying that the suggested firing aspects of Purkinje cells and cerebellar nuclei can be obtained, the present results can be corrected quantitatively and proposed as more realistic findings.

## Conclusions

Present simulation reproduced the sustained firing of the cerebellar nuclei during motor learning in which the climbing fiber inputs to purkinje cell are continuously randomized, and revealed a suggestive factor regarding the cerebellar neural mechanism of motor-learning-induced task-specific sustained tremor, which is like a symptom of FTSD. Based on the results of this study, we also proposed a motor learning paradigm, “undesired motor learning”, in which the motor goal is too minute to be achieved by repetition alone and converges to a different outcome than the desired one. This finding would be verified as the principle of FTSD acquisition by motor learning through learning experiments using actual organisms and will be expected to contribute to the prevention and treatment of the disease as well as to the consideration of motor learning strategies for expert athletes or musicians.

## Acknowledgments

We would also like to thank Leave a Nest Co., Ltd. for funding this research.

